# Evidence for an Integrated Bilingual Language System from Discourse Tasks in Aphasia

**DOI:** 10.64898/2025.12.17.694682

**Authors:** Xuanyi Jessica Chen, Manuel J. Marte, Swathi Kiran, Esti Blanco-Elorrieta

## Abstract

**Background:** The extent to which bilingual individuals represent and process their two languages within a shared or partially distinct neural architecture remains a topic of ongoing debate. While both parallel and divergent patterns of impairment have been reported in bilingual aphasia, such findings likely reflect a spectrum of representational overlap influenced by dominance, proficiency, and task demands. Critically, few studies have examined how breakdown manifests across multiple levels of linguistic structure using ecologically valid, discourse-based tasks.

**Aims:** This study investigates whether Spanish-English bilinguals with aphasia exhibit parallel or dissociable patterns of impairment across their two languages, focusing on naturalistic narrative production and fine-grained analysis of error types and code-switching.

**Methods & Procedures:** Thirteen bilingual individuals with aphasia following acquired brain injury produced story retellings in both languages. Speech samples were transcribed and coded for phonological, morphological, syntactic, and semantic errors, and for the word type at which they occurred. Code-switches were also identified and categorized. Analyses included generalized linear modeling and unsupervised clustering.

**Outcomes & Results:** While participants made more errors in their non-dominant language, the structure and distribution of errors were highly similar across languages. Clustering algorithms revealed that impairments were parallel across languages and did not group by language dominance. Code-switching occurred more frequently from the non-dominant to the dominant language, consistent with activation-based lexical selection.

**Conclusions:** Findings support an integrated bilingual language system that spans multiple levels of linguistic representation, modulated by language dominance. Naturalistic discourse tasks allow for richer characterization of bilingual language breakdown and may better inform both theoretical models and clinical management of bilingual aphasia.

## 1 Introduction

Understanding how bilingual individuals process and represent their two languages remains a central question in cognitive neuroscience, especially in the context of brain injury, where disruption to the language system offers a unique opportunity to infer its underlying architecture. At the heart of this inquiry lies a fundamental theoretical question: do bilinguals represent and process both languages within a single, unified neurocognitive architecture, or do they rely on partly distinct systems? A fully integrated model assumes that once a neural mechanism exists for a given linguistic operation (e.g., plural inflection), it serves that function across languages, treating cross-language variation similarly to within-language variation (e.g., across registers or dialects; Blanco-Elorrieta & Caramazza, 2021). In contrast, other perspectives argue that features such as formal grammatical structure (MacWhinney et al., 2005; MacWhinney, 2022), age of acquisition (Singleton, 2005; Singleton & Leśniewska, 2021), and usage context can give rise to at least partially distinct representations and processing routines.

These views are not mutually exclusive, and growing evidence suggests that the bilingual mind exhibits both integration and separation depending on the linguistic domain and level of representation (Abutalebi & Green, 2013; Van de Putte et al., 2017, 2018; Malik-Moraleda et al., 2024). The present study directly tests these competing accounts by examining how bilingual language breakdown unfolds across multiple linguistic levels following brain injury.

Specifically, we investigate whether Spanish–English bilinguals with post-stroke aphasia show parallel or dissociable patterns of impairment across their two languages, and whether code-switching behavior provides converging evidence for an integrated or partially distinct language system. By analyzing naturalistic, discourse-based production, we aim to capture how bilingual language organization manifests in people with aphasia under real-world communicative conditions rather than decontextualized laboratory tasks.

Within healthy populations, much prior work has focused on the lexico-semantic level (Chee et al., 1999; Dehaene et al., 1997; Hasegawa et al., 2002; Yang et al., 2017; Molinaro et al., 2024), with relatively limited attention to morphosyntactic and phonological representations (Declerck et al., 2020; Ou et al., 2020). While conceptual representations are generally believed to be shared across languages (Van de Putte et al., 2018; Chen et al., 2024), debates continue over whether morphology, syntax, and phonology are supported by common or distinct neural mechanisms. Findings vary from overlapping activation patterns for L1 and L2 (Tettamanti et al., 2002; Musso et al., 2003; Sakai et al., 2004; Perani & Abutalebi, 2005; Yang et al., 2017) to evidence of differential activity for L1 and L2 in distinct brain regions (Lehtonen et al., 2009; Golestani et al., 2006; Ip et al., 2017), and even language-specific differentiation across regions of interest (Gao et al., 2023; Meykadeh et al., 2021; see Cargnelutti et al., 2019 for meta-analysis; Biondo et al., 2023 for review).

Yet most of this evidence is correlational, leaving open the question of causality: whether the same neural substrates are not only co-activated across languages but also jointly necessary for their function. Studying bilinguals with post-stroke aphasia provides a rare opportunity to test this directly: lesions to shared neural resources should lead to parallel impairments in both languages, whereas lesions affecting language-specific regions should yield selective deficits in one language only. In this way, bilingual aphasia serves as a natural lesion model that can arbitrate between integrated and partially distinct accounts of bilingual language representation. By analyzing the nature and distribution of errors across linguistic levels, we can infer whether both languages rely on common processing mechanisms or on separable neural subsystems.

In addition to linguistic level, word class provides a finer-grained window into the organization of bilingual language representations. A large body of research has shown that aphasia can differentially affect content words (nouns, verbs, adjectives) and function words (articles, prepositions, pronouns, conjunctions), reflecting their distinct roles in language production and neural representation. Content words carry rich semantic and conceptual information, whereas function words primarily encode grammatical relations and structural dependencies (Friederici, 2011; Pulvermüller, 2002). These categories are supported by partially separable neural substrates: content words engaging temporo-parietal semantic regions and function words relying more heavily on left inferior frontal and temporal areas involved in morphosyntactic processing (Hagoort, 2005; Tyler et al., 2011).

Clinically, this distinction is also evident: agrammatic and non-fluent aphasias often involve disproportionate difficulty with function words and inflectional morphology, while fluent aphasias tend to spare grammatical morphemes but produce semantically empty content (Caramazza & Zurif, 1976; Bastiaanse & Thompson, 2012). Because bilingual aphasia might manifest either as parallel or dissociable breakdown across these lexical categories, analyzing errors by word class offers a more precise diagnostic lens on the structure of the bilingual language system. If both languages share a unified representational and control architecture, we would expect similar function–content asymmetries in both languages; if they are partly distinct, the relative vulnerability of these categories could diverge. Incorporating word class thus allows us to examine not only whether the two languages break down in parallel, but how deeply that parallelism extends into the grammatical versus lexical strata of language.

Foundational neuropsychological work on bilingual aphasia has documented both parallel and differential patterns of post-stroke impairment, and these contrasting outcomes have been central to the debate over whether bilingual language is supported by shared or partially distinct systems. Classic reports by Paradis (2001) and by Fabbro (2001) described cases in which one language was disproportionately or selectively impaired, in some instances with marked dissociations at specific linguistic levels (e.g., lexical access impaired in one language but preserved in the other, or grammatical morphology disrupted in only one language), and in some cases with asymmetric recovery trajectories across languages over time. Such cases have been interpreted as evidence that, at least under some lesion and experience profiles, the two languages can rely on partly independent neural resources and/or processing routines. In contrast, group-level work has mainly found broadly parallel impairment across languages in bilingual persons with aphasia, such that both languages show qualitatively similar error types and breakdown patterns, even when they differ in overall severity (e.g., more errors in the less proficient or less frequently used language; Goral et al., 2006; Green et al., 2010; Kohnert, 2004; Kiran & Iakupova, 2011). On this view, apparent between-language differences largely reflect graded accessibility (frequency, dominance, pre-morbid use) rather than distinct representational architectures. Together, these literatures suggest that bilingual aphasia outcomes span a continuum from tightly coupled to partially dissociable, and that this continuum is influenced by factors such as age of acquisition, dominance, lesion site, and post-stroke language use (Kiran et al., 2013).

Despite the richness of this work, two key gaps remain and directly motivate the present study. First, most prior studies have focused on a narrow set of tasks (e.g., picture naming, translation, word-level repetition) and therefore speak primarily to lexico-semantic retrieval.

Much less is known about whether bilinguals show parallel versus dissociable impairment profiles across multiple levels of linguistic representation (i.e., phonology, morphology, syntax, and semantics) when producing connected speech.

Second, relatively little work has examined code-switching in spontaneous discourse as a theoretically meaningful diagnostic behavior. Code-switching is often described clinically as a symptom of impaired control, yet it may instead reflect adaptive recruitment of whichever lexical or structural resource is most available at a given moment. Critically, this behavior provides a dynamic window into the functional organization of the bilingual system and therefore bears directly on the question of whether both languages share or segregate neural resources. If both languages draw on shared neural substrates, switches should emerge as fluid transitions within a single integrated network rather than as costly shifts between distinct systems. Under this activation-based view, switches are expected to occur more frequently from the non-dominant to the dominant language (i.e., the language with overall greater activation; Blanco-Elorrieta & Caramazza, 2021). In contrast, partially separated or inhibitory models (Green, 1998) propose that producing the non-dominant language requires active suppression of the dominant one, such that failures of inhibition result in “reverse dominance,” i.e., more frequent switching into the non-dominant language (Gollan et al., 2014; Li & Gollan, 2018; Fadlon et al., 2019). Examining the direction and frequency of code-switches in bilingual PWA thus provides a behavioral proxy for how tightly coupled or segregated their two languages are at the neural level. The current study addresses these gaps by analyzing naturalistic narrative production in Spanish–English bilinguals with aphasia, coding error types across multiple representational levels and examining the directionality and function of spontaneous code-switches.

Language typology may also play a role in shaping bilingual language organization. Studies comparing typologically distant language pairs (e.g., English–Mandarin, Spanish–Arabic) suggest that linguistic distance can influence cross-language transfer, representational overlap, and recovery profiles (Ghazi-Saidi & Ansaldo, 2017). That said, if the principles of language representation are indeed universal, as posited by many neurocognitive models, then typological distance alone should not necessarily result in fundamentally different neural architectures. At present, no clearly articulated theory specifies how or why structural differences in morphology, syntax, or phonology would causally give rise to distinct neural or cognitive organizations. This remains an important area for future theoretical development. The current study focuses on Spanish and English, two Indo-European languages that, while different in grammatical and phonological properties, share structural similarities that may facilitate integration (e.g., subject-verb-object word order, alphabetic script).

Crucially, regardless of typological distance, different theoretical models make distinct predictions about how linguistic deficits and code-switching behaviors (i.e., alternating between languages within a conversation or utterance) should appear in bilingual PWA. Integrated models (e.g., McClelland & Rumelhart, 1981; Rumelhart & McClelland, 1982; Blanco-Elorrieta & Caramazza, 2021) predict that an impairment at a given representational level should affect both languages similarly in quality, with differences arising only in degree (for example, greater impairment in the non-dominant language due to less frequent use; Nadeau, 2019, or lower proficiency Peñaloza et al., 2020a). Under this view, parallel error profiles would be expected across languages, even if severity varies (Jacquemot et al., 2007; Jeffries et al., 2006; Lallini et al., 2007; Martin et al., 1996). Crucially, these predictions must be evaluated separately for each linguistic level: it is entirely plausible that phonological representations might be integrated while morphosyntactic representations remain distinct.

In contrast, partially separated models predict that impairments can show language-specific dissociations—such as impaired phonology in one language but intact phonology in the other, or distinct grammatical error profiles (MacWhinney, 2022; Paradis, 2004). Crucially, such dissociations have been observed in single-case reports (e.g., Fabbro, 2001; Paradis, 2001). Thus, our aim here is to test the extent to which integration holds across linguistic levels using naturalistic production tasks.

In all, the hypotheses we test are as follows.

1. If representations are fully integrated, error patterns should be qualitatively identical across languages, differing only in quantity based on dominance or proficiency.
2. If representations are partially separated, we should observe Language × Linguistic Level interactions, with some levels showing parallel deficits and others showing dissociations.
3. If activation-based lexical selection governs bilingual production, code-switching should predominantly occur from the non-dominant to dominant language.

In the current study, we aim to characterize bilingual aphasia by analyzing the typology of language impairments and the direction and function of code-switches during narrative production. Rather than relying on decontextualized tasks (e.g., picture naming), we use open-ended storytelling to elicit discourse that spans multiple representational levels and allows us to examine real-time language use across both languages. This approach provides an ecologically valid and theoretically rich opportunity to trace patterns of impairment and code-switching across specific linguistic domains. By doing so, we aim to highlight possible areas of integration and dissociation, and consider how these patterns might inform models of bilingual language organization.

## 2 Methods

### 2.1 Participants and Experimental procedure

We collected data from 13 English-Spanish bilinguals with aphasia (9 F, 4 M; age 26-68; *M* = 49, *SD* = 13.74). All PWA were living in the United States at the time of the study. Their language profile was assessed with the Language Use Questionnaire (LUQ; Kastenbaum et al., 2019; Marte & Carpenter et al., 2022), a notably comprehensive test that measures daily language use, language proficiency of family members, language of instruction at each level of education, language exposure, language confidence and self-reported language ability in hearing, speaking, reading and writing across their lifetime (see Fig. 1A top panel).

**Figure 1.**
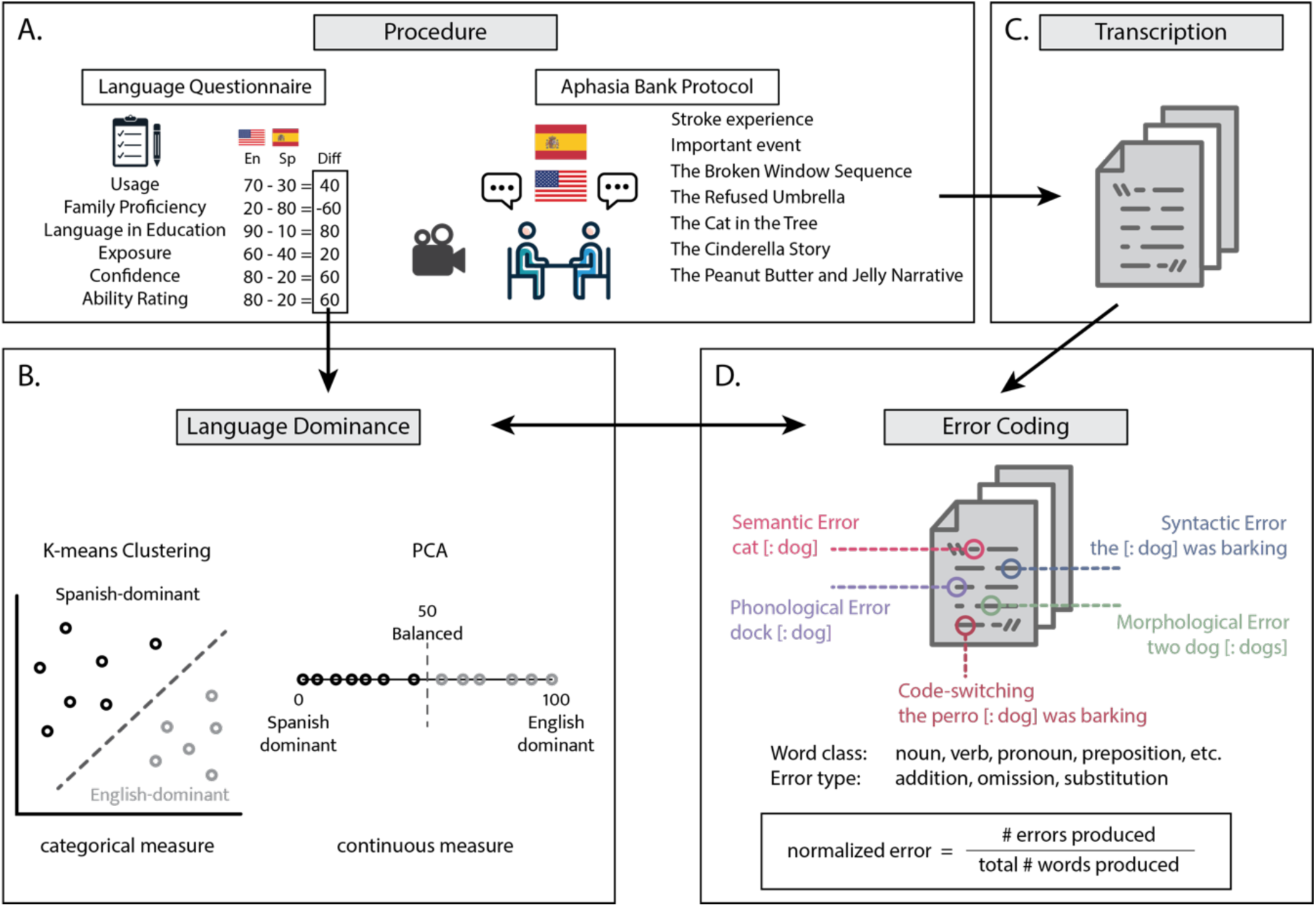
**A. Testing and analyses procedure**. During the testing session, persons with aphasia (PWA) completed the Language Use Questionnaire (LUQ; Kastenbaum et al., 2019) and the Aphasia Bank Protocol (MacWhinney et al., 2011) in English and Spanish**. B. Language dominance calculation**. The difference between languages in every measure of the language questionnaire was calculated and used to derive two measures for language dominance: a categorical measure of language dominance using K-means clustering and a continuous measure using PCA**. C. Transcription**. PWAs’ speech was transcribed by two bilingual native speakers. **D. Error Coding.** PWAs’ speech was coded for errors and code-switches. For each error, we coded for the level of representation (semantic, syntactic, phonological, morphological), word class (noun, verb, adjective, adverb, pronoun, determiner, preposition, conjunction), and error type (addition, omission, substitution). Normalized errors were calculated by scaling the raw frequency of errors by the total number of words produced (excluding repetition and retraction).

All PWAs were tested during the subacute and chronic phase of their stroke and completed the Aphasia Bank protocol (MacWhinney et al., 2011), which aims to elicit PWA’s speech by asking them to: i) describe their stroke experience and a memorable event, ii) describe some pictures (the Broken Window sequence, the Refused Umbrella, the Cat in the Tree), iii) tell the Cinderella story, and iv) describe the steps required to make a Peanut Butter and Jelly sandwich. All testing was administered by trained, bilingual Spanish-English speech-language pathologists and research assistants. Each testing session was conducted entirely in one language (either English or Spanish), with examiners using only the target language during task administration to maintain consistent language context. Testing languages were counterbalanced across sessions, and examiners were instructed to minimize verbal feedback during narrative production to avoid influencing participants’ spontaneous language production and code-switching behaviors. The speech output of these tasks constitutes the corpus for our analyses. Ten patients had confirmed left-hemisphere lesions, while three had unspecified lesion lateralization in their medical records. Etiologies included ischemic stroke (n = 5), hemorrhagic stroke (n= 1), unspecified CVA (n = 5), Moyamoya disease (n = 1), and brain tumor (n = 1). The Western Aphasia Battery (WAB; Kertesz, 2007) was additionally completed to determine the severity of the aphasia in both languages. All PWAs received a WAB aphasia quotient score of ≥60 in both languages, indicating that they were mild or moderate aphasia per WAB classification (see Table. 1). WAB testing was counterbalanced across languages and sessions to control for order effects. All tests were administered by a trained speech-language pathologist either in person or on zoom.

### 2.2 Language dominance calculation

Critical to our hypothesis testing is to determine i) whether there is a *quantitative* difference across languages that reflects higher resilience to damage of higher frequency elements (here understood also as dominant language elements), and ii) whether a *qualitatively* different profile of errors emerges between dominant and non-dominant language. Both of these require defining each PWA’s pre-morbid language dominance, which is non-trivial due to the varying nature of every person’s language experience.

Traditionally, AoA and/or proficiency have been used to operationalize language dominance, but they are too one-dimensional to to capture the multidimensional nature of language experience. To address this, we adopted a comprehensive, data-driven approach by incorporating all dimensions from the Language Use Questionnaire (LUQ; Kastenbaum et al., 2019; Marte & Carpenter et al., 2022)—including language usage, exposure, proficiency of family members, educational language history, confidence, and self-rated ability across modalities. We calculated the difference between English and Spanish for each dimension, and then applied both K-means clustering and Principal Component Analysis (PCA) to these difference scores to classify participants’ relative language dominance (Fig. 1B).

K-means clustering is an algorithm that takes multidimensional data (in our case, the data from every language experience dimension for each subject) and groups it on K number of clusters based on similarity across all dimensions. Here, we used the algorithm to divide the data into two clusters: Spanish-dominant and English-dominant. The decision to apply a two-cluster (K=2) K-means solution was theoretically motivated. Our core predictions center on asymmetries between a speaker’s dominant and non-dominant language, predictions that hold even when the dominance is subtle. Unless a participant is exactly balanced, even small asymmetries in language experience are theoretically meaningful and may shape processing outcomes. In fact, including participants with marginal dominance differences provides a more stringent test of our hypotheses. If effects emerge across this full range, they are less likely to be artifacts of selecting only clearly unbalanced individuals and more likely to reflect generalizable patterns of bilingual processing.

K-means clustering grouped participants into two data-driven categories. The algorithm classified 7 participants in the Spanish-dominant group and 6 in the English-dominant group, based on their composite language profiles.

To preserve the continuity of dominance and to validate the clustering outcome, we also applied PCA, which reduces multidimensional data into a continuous scale. PCA is a linear dimensionality reduction technique, which takes high-dimensional data (i.e., the data from every dimension in the language experience questionnaire) and transforms it into low-dimensional data by minimizing redundant information across related measures. In our case, we want to obtain one variable from the language experience questionnaire that reflects subjects’ relative language dominance. We first calculated the difference between English and Spanish in each of the language experience dimensions (e.g., proficiency), and submitted the vector containing these difference scores to a PCA analysis. Then, we took the first component, which is the component that best explains the variance in the data (in our case, 88.86%), and we obtained a raw language dominance measure by calculating the projection of all dimensions for each subject along this first component. To determine the directionality of this measure (i.e. whether positive indicates English-dominant or Spanish-dominant), we inspected the language profile of the two subjects who received the maximum and minimum values. Finally, we scaled and recentered the raw measures to the range of 0-100, such that the most Spanish-dominant measure is mapped to 0 and the most English-dominant measure is mapped to 100. The value represents the relative dominance of English or Spanish, based not just on Age of Acquisition or Proficiency, but rather based on the compound of all the answers they provided in the comprehensive Language Use Questionnaire (LUQ; Kastenbaum et al., 2019).

Importantly, the results across the categorical and continuous measures in our analysis were fully consistent: participants with a score of 50 or more in PCA were clustered in the English-dominant cluster by K-means, and those with a score of less than 50 clustered as Spanish-dominant. This strong convergence across methods enhances confidence in our classification scheme, while also preserving the ability to explore graded dominance effects. This dual approach ensures both theoretical precision and empirical robustness in our treatment of language dominance.

### 2.3 Data transcription and coding

Recordings of the testing sessions were transcribed in the Computerized Language Analysis Software (CLAN; MacWhinney, 2000; see Fig. 1 for a detailed description of the procedure). All recordings were independently transcribed and coded following the AphasiaBank transcription guidelines by native speakers of each language. To assess inter-rater reliability, we followed Koo and Li’s (2016) guidelines for selecting and reporting intraclass correlation coefficients (ICC). Specifically, we calculated a two-way random-effects, absolute-agreement, single-measure ICC, which is appropriate when each subject (here, each transcript) is independently rated by two coders randomly selected from a larger pool. ICC values were computed for both transcription accuracy and error categorization across all linguistic levels (phonological, morphological, syntactic, and lexico-semantic). According to Koo and Li (2016), ICC values below .50 indicate poor reliability, between .50–.75 moderate, between .75–.90 good, and above .90 excellent reliability. Prior to formal analysis, coders independently transcribed and coded 20% of the dataset (randomly selected across participants and languages). ICCs were then computed for (a) number of words transcribed, (b) number of coded errors per linguistic level, and (c) classification consistency of error types. All reliability estimates exceeded the threshold for *good* reliability (ICC > .80).

To the extent that AphasiaBank was created for English, the range of error codes did not cover all the possible errors one could encounter in Spanish. Thus, we developed and expanded this guide to incorporate all the types of mistakes PWAs could make in Spanish. We have made the guide including all the possible error types and examples in English and Spanish publicly available in OSF: https://osf.io/6mwyx/?view_only=ebbfad1c0e0148c0aad2c473c4ea51f9).

The transcribers coded each error for:

1. The linguistic level where they occurred (i.e., syntactic, semantic, phonological, and morphological).

a. We coded as syntactic errors occasions when required components of a sentence were missing (e.g., “the [dog] was barking”), duplicated e.g., “the dog dog was barking”), or produced in a wrong order (e.g., “the was dog barking”).
b. We coded as lexico-semantic errors both when PWAs substituted a target word for a semantically related one, and when they had word finding problems that resulted in no production.
c. Phonological and morphological errors included occasions where PWAs added, omitted, or produced an incorrect phoneme/morpheme.
2. The class of the target word (e.g. noun, verb, pronoun, preposition, adjective, adverb).
3. The type of error the PWAs made (i.e., addition, substitution, omission; see Fig. 1 bottom right).

When a production contained multiple errors, we coded them all. For example, when a PWA produced ‘daufter’ for the target word ‘son’, it was coded as a lexico-semantic error (as they tried to produce the semantically related word ‘daughter’ for ‘son’) and a phonological error (for the additional phoneme /f/).

We also identified where production switched from one language to another (i.e., there was a code-switch) and coded for the class of the code-switched word (e.g., noun, verb…). When the code-switch occurred for multiple-word phrases, we indicated both that the switch happened at the phrase level and the class of the first word of the phrase.

A potential issue with using the raw number of errors/code-switches in spontaneous speech production is the considerable variation in the amount of speech produced by each PWA. It is crucial to address this because finding a mistake every 1000 words is significantly different from finding one in every other word. Therefore, to obtain a meaningful measure of PWA’s impairments, it is necessary to normalize the error rates. Two common methods of normalization in the literature involve dividing the raw number of errors by the total number of words or by the total number of utterances. For people with aphasia, the total number of utterances is often inflated by short discourse exchanges (e.g., “yeah,” “mm-hmm”), so we chose to normalize by the total number of words instead. Given that repetitions and reiterations common in PWA might also inflate the word count, we settled for normalizing the errors using the total number of words in that language excluding repetitions and retractions as the denominator. Ultimately, the three measures of discourse production (total number of utterances, and total number of words with and without retractions) were highly correlated (*r* > 0.9; *p* < 0.001), such that the choice of denominator did not have significantly impact the normalization (Fig. 1D). We identified and excluded outliers in our normalized measures using the maximum normalized residual test (Grubbs’ test), which led to the exclusion of one participant for the code-switching analyses.

### 2.4 Statistical analyses

To explore the impact of premorbid language dominance on post-stroke linguistic abilities, we calculated the correlation between language dominance (as measured by PCA) and both (i) the WAB aphasia severity score and (ii) the normalized amount of errors produced. Based on previous research (Kuzmina et al., 2019; Peñaloza et al., 2020b), we expected the dominant language to show milder impairment, reflected in higher WAB scores and fewer errors.

To examine how error rates varied as a function of linguistic and lexical factors, we fitted linear mixed-effects (LME) models with normalized error rate as the dependent variable and Language Dominance (dominant vs. non-dominant), Level of Representation (phonological, morphological, syntactic, lexico-semantic), and Word Class (noun, verb, adjective, adverb, pronoun, determiner, preposition, conjunction) as fixed effects. Subject was included as a random intercept.

We followed a two-step analytic procedure:

1. Model selection: Candidate models were compared using the Bayesian Information Criterion (BIC) to identify the most parsimonious model balancing explanatory power and complexity (Burnham & Anderson, 2004). This procedure was used exclusively to determine model structure, not to test significance.
2. Significance testing: After identifying the optimal model, we assessed the significance of fixed effects and interactions within that model using Type III F-tests with Satterthwaite approximation for degrees of freedom (via the *lmerTest* package).

The final model that minimized the BIC included the main effects of Language Dominance, Level of Representation, and Word Class, as well as the Level of Representation × Word Class interaction. Adding any interaction terms involving Language Dominance increased the BIC by more than 30 points, providing strong evidence that those terms did not improve model fit.

Finally, we also analyzed the effect of Language Dominance on the amount of code-switching PWAs produced. To this end, we calculated the correlation between PCA language dominance and the normalized amount of code-switches. We further fitted an LME model to evaluate how Language Dominance and Word Class influence the normalized code-switching measure (random effect: subject). Both models predict a significant effect of language dominance on the amount of code-switches produced, but in opposite directions. Under an integrated language system, where lexical selection occurs on the bases of highest level of activation (Blanco-Elorrieta & Caramazza, 2021), code-switches should be more prevalent from the non-dominant to the dominant language, because items in the dominant language have higher baseline activation and are less likely to be impaired after brain damage. Contrastingly, under inhibition-based lexical selection approaches, one would expect a ‘reverse dominance’ effect: more switches from the dominant language to the non-dominant language (Gollan et al., 2014; Li & Gollan, 2018; Fadlon et al., 2019), as a consequence of the hypothesized harder inhibition required to suppress elements from the dominant as compared to the non-dominant language.

## 3 Results

**All effects reported below are derived from the final LME model described in Section 2.4.**

### 3.1 Uneven quantitative loss across languages: dominance predicts error quantity

PWAs’ post-stroke language ability was determined by their premorbid language profile as shown by a significant correlation between the Language Dominance score (PCA) and i) the difference in the WAB score (Table 1; *r*=0.83, *p*<0.001; Fig. 2A) and ii) the amount of errors produced in the AphasiaBank tasks (Fig. 2B; *r*=-0.68, *p*=0.01). As predicted, the more dominant language was more spared in our bilingual PWAs. This finding was confirmed by the LME analysis, which uncovered a main effect of Language Dominance (*F*=14.32, *p*<0.001), whereby PWAs produced more errors in their non-dominant language than in their dominant language across all levels of representation and word classes (Figure 2C-D, Figure 3A-B).

**Figure 2.**
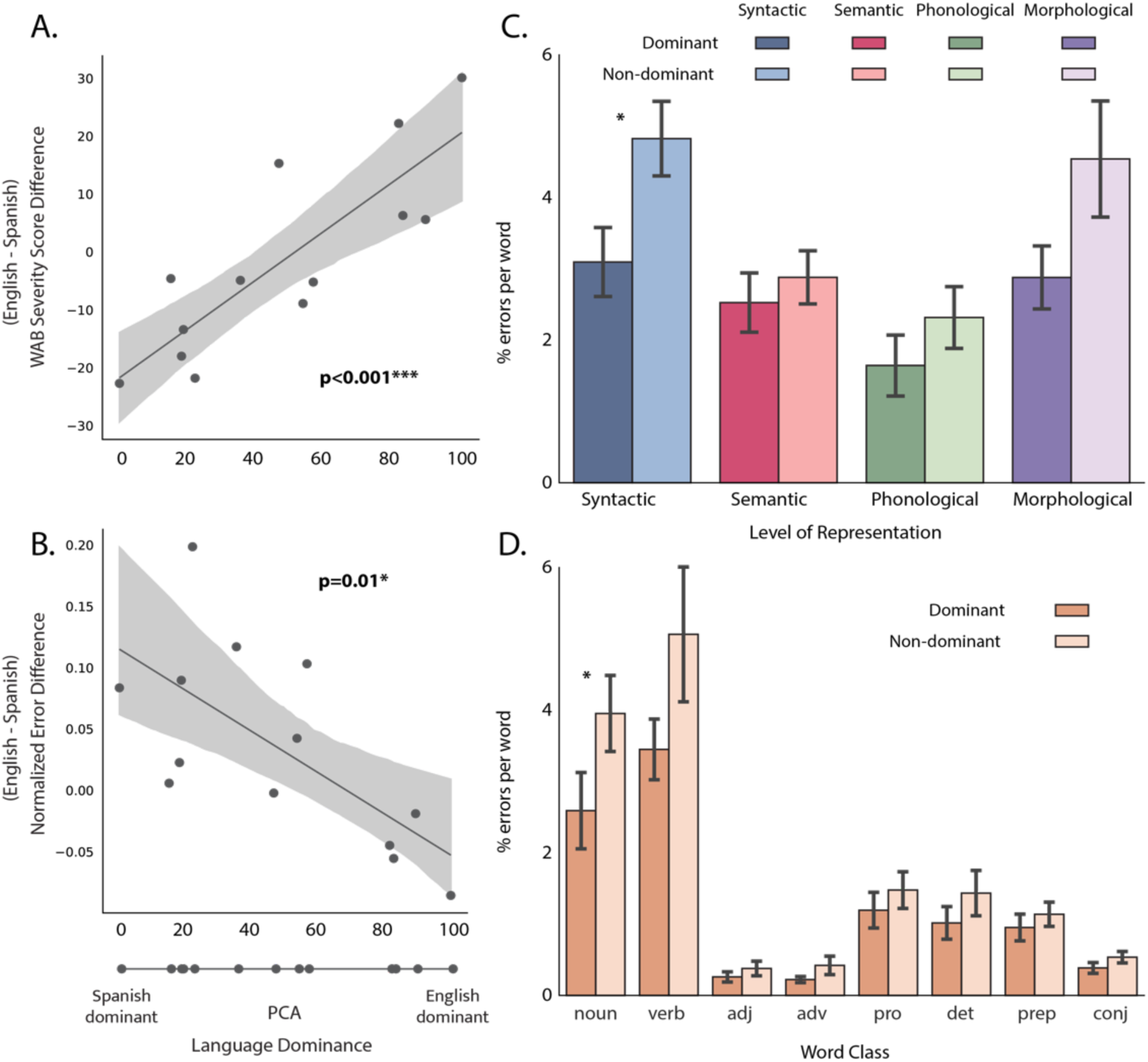
Effects of language dominance on post-stroke language abilities. A) shows the relationship between language dominance and difference in WAB severity score. B) shows the relationship between Language Dominance and amount of produced errors. Panels C and D show the error rates at each level of representation and word class in the dominant and non-dominant language, showing equal patterns of impairments across languages.

**Figure 3.**
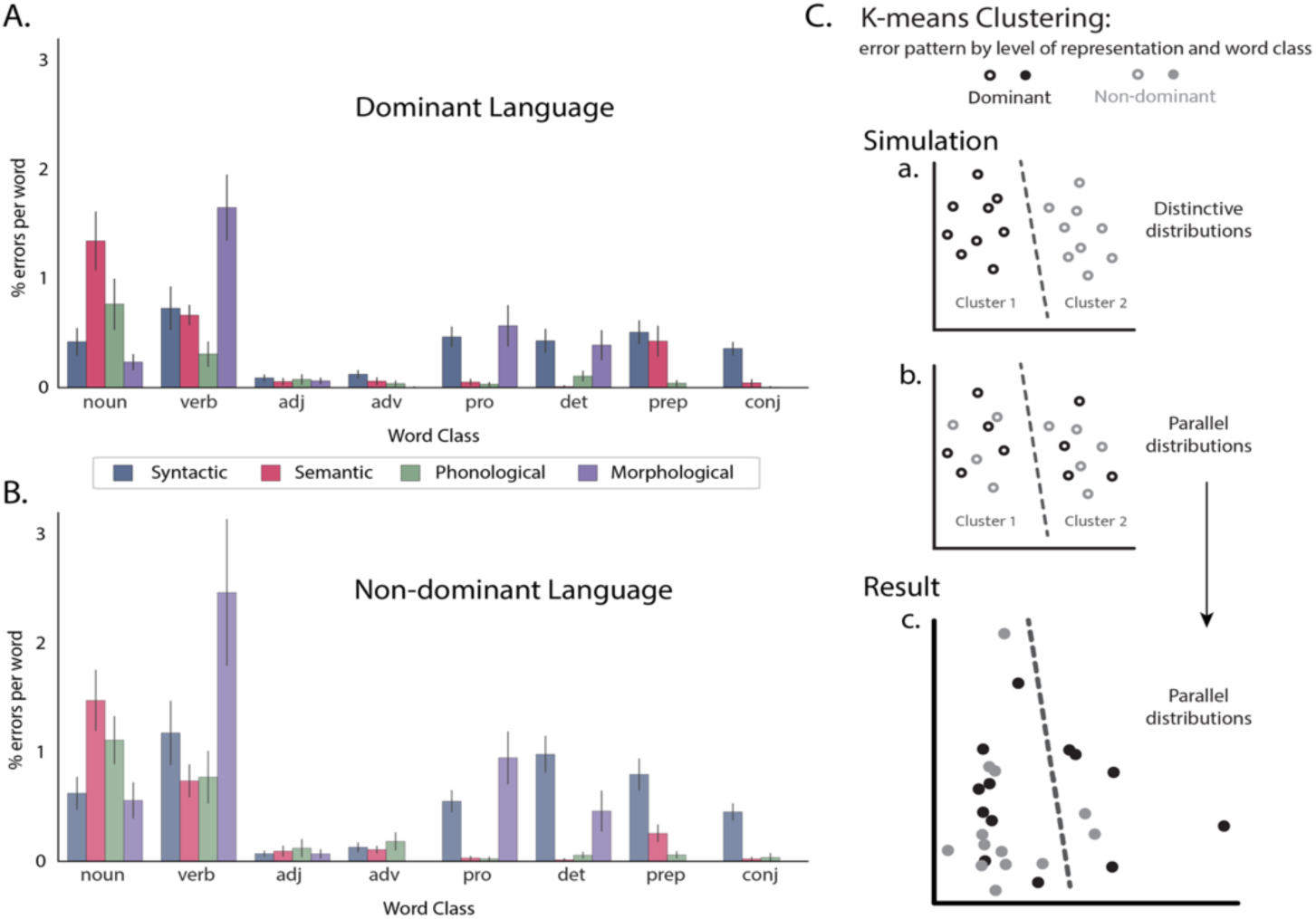
Effect of language dominance on error distribution. A-B: error distribution across level of representation and word class in dominant and non-dominant language. C: K-means clustering of error pattern by level of representation and word class in simulated data (a-b) and actual result (c).

**Table 1.**
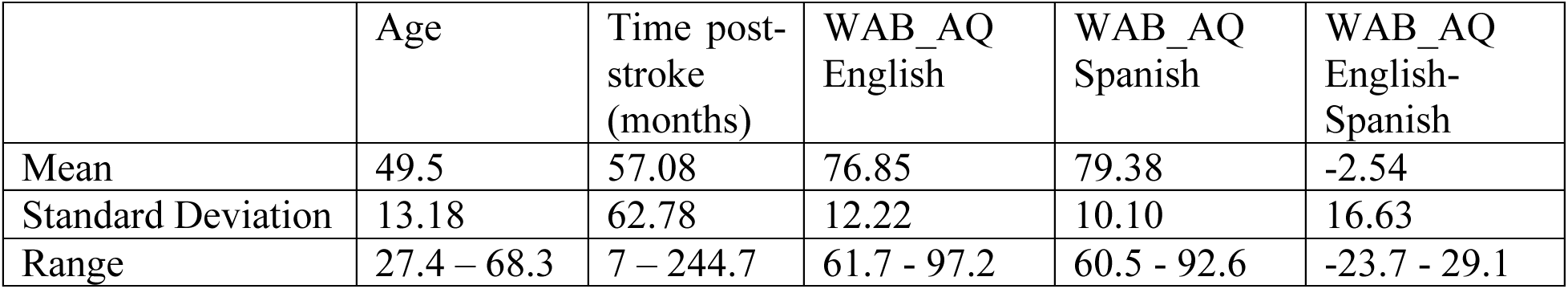
Demographics and clinical characteristics of the PWA (n = 13). Values include time post-stroke (months) and Western Aphasia Battery-Revised Aphasia Quotient (WAB AQ) scores in both English and Spanish, with the difference between languages (English minus Spanish).

|  | Age | Time post-stroke (months) | WAB_AQ English | WAB_AQ Spanish | WAB_AQ English-Spanish |
| --- | --- | --- | --- | --- | --- |
| Mean | 49.5 | 57.08 | 76.85 | 79.38 | -2.54 |
| Standard Deviation | 13.18 | 62.78 | 12.22 | 10.10 | 16.63 |
| Range | 27.4 – 68.3 | 7 – 244.7 | 61.7 - 97.2 | 60.5 - 92.6 | -23.7 - 29.1 |

### 3.2 Parallel qualitative loss across languages: dominance does not determine error typology

To examine whether dominant and non-dominant languages show qualitatively parallel error patterns (supporting a unified language system for bilingual individuals) or whether there are selective deficits that manifest solely in one language (revealing that languages operate on independently represented systems), we focused on evaluating the interaction terms between Language Dominance, Word Class, and Level of Representation.

The final LME model (see Section 2.4) included the main effects of Language Dominance, Level of Representation, and Word Class, plus the Level × Word Class interaction. Model comparison using BIC confirmed that adding either a Dominance × Level or Dominance × Word Class interaction worsened model fit (ΔBIC = +31.03). According to conventional thresholds (Raftery, 1995), a BIC increase greater than 10 constitutes *very strong* evidence against the more complex model. In other words, the inclusion of Dominance × Level or Dominance × Word Class interactions did not improve the model’s explanatory power sufficiently to offset their added complexity, indicating that these interactions are not supported by the data. Conceptually, this outcome is equivalent to finding that those interaction effects are not statistically significant (Burnham, & Anderson, 2004; Wagenmakers, & Farrell, 2004). In short, language loss shows a parallel qualitative profile across dominant and non-dominant languages (see Fig. 3A–B).

Within the final model, we observed a main effect of Level of Representation (*F* = 11.31, *p* < .001), driven by higher error rates in morphology and syntax than in lexico-semantic or phonological levels. There was also a main effect of Word Class (*F* = 51.90, *p* < .001), with most errors occurring in nouns and verbs. A significant Level × Word Class interaction (*F* = 12.97, *p* < .001) reflected that different word types afford distinct opportunities for errors (e.g., semantic errors are more frequent in content words, whereas function words are more constrained).

To further confirm that language dominance does not introduce qualitative divergence in error profiles, we conducted an unsupervised K-means clustering analysis as a complementary, data-driven test. We generated 26 vectors (one per participant per language), each containing 32 dimensions (4 representational levels × 8 word classes), capturing the error distribution for that language. We then asked whether the algorithm could recover two clusters corresponding to dominance status. If dominance meaningfully shaped patients’ impairment profile, the algorithm should separate the data into distinct clusters for dominant versus non-dominant languages; if impairment is parallel across languages, the clustering should be blind to language dominance and grouping should not be based on this dimension. This logic is illustrated in Figure 3C, panels a and b, which simulate the two possible outcomes: (a) distinctive clustering by language, and (b) parallel distributions across languages.

The actual result of our clustering analysis is shown in Figure 3C, panel c. The algorithm returned two clusters of similar size, but each contained a mix of dominant and non-dominant language data points (Cluster A: 8 dominant / 11 non-dominant; Cluster B: 5 dominant / 2 non-dominant). To quantify statistically how well the clustering aligned with the grouping expected from dominance, we used the Adjusted Rand Index (ARI). The ARI measures the similarity between two groupings on a scale from –1 to 1, where 1 indicates a perfect match, 0 indicates similarity no better than random chance, and negative values indicate systematic disagreement. Here, we compared our grouping to the language-dominance based grouping and the observed ARI was 0.02, effectively at chance, showing that the clustering bore no relationship to what would be expected if dominance shaped the error patterns.

To further complement the unsupervised K-means analysis, we also tested whether a supervised model (i.e., one that is explicitly told which samples are dominant vs. non-dominant) could learn to classify the error patterns by dominance. We used a support vector machine (SVM), a standard classifier in machine learning, and evaluated it using 13-fold cross-validation so that each prediction was made on data the model had not seen during training. The model’s accuracy was 42%, below chance level (50%) when distinguishing two groups. This result shows that even when given the correct labels to learn from, the model could not find dominance-related structure in the data. Together, these results support that “Language Dominance” was not a meaningful dimension for grouping the data, supporting the conclusion that error distributions across languages were qualitatively parallel.

### 3.3 Uneven code-switches across languages: dominance predicts language switch quantity and directionality

Consistent with the prediction of an activation-based integrated model (Blanco-Elorrieta & Caramazza, 2021), we observed more code-switches from the non-dominant to the dominant language than vice versa (Fig. 4). We found a significant correlation between Language Dominance and the difference in the normalized code-switching measure (*r*=-0.64, *p*=0.02) and a main effect of Language Dominance in the LME model (*F*=5.33, *p*=0.02). There was no significant main effect of Word Class (*F*=1.85, *p*=0.08), and we did not include interaction terms as the model without them received the lowest BIC scores (-1572 vs -1549; D = 23.75), suggesting that including interaction terms did not provide a better estimation of the data over and beyond the added model complexity.

**Figure 4.**
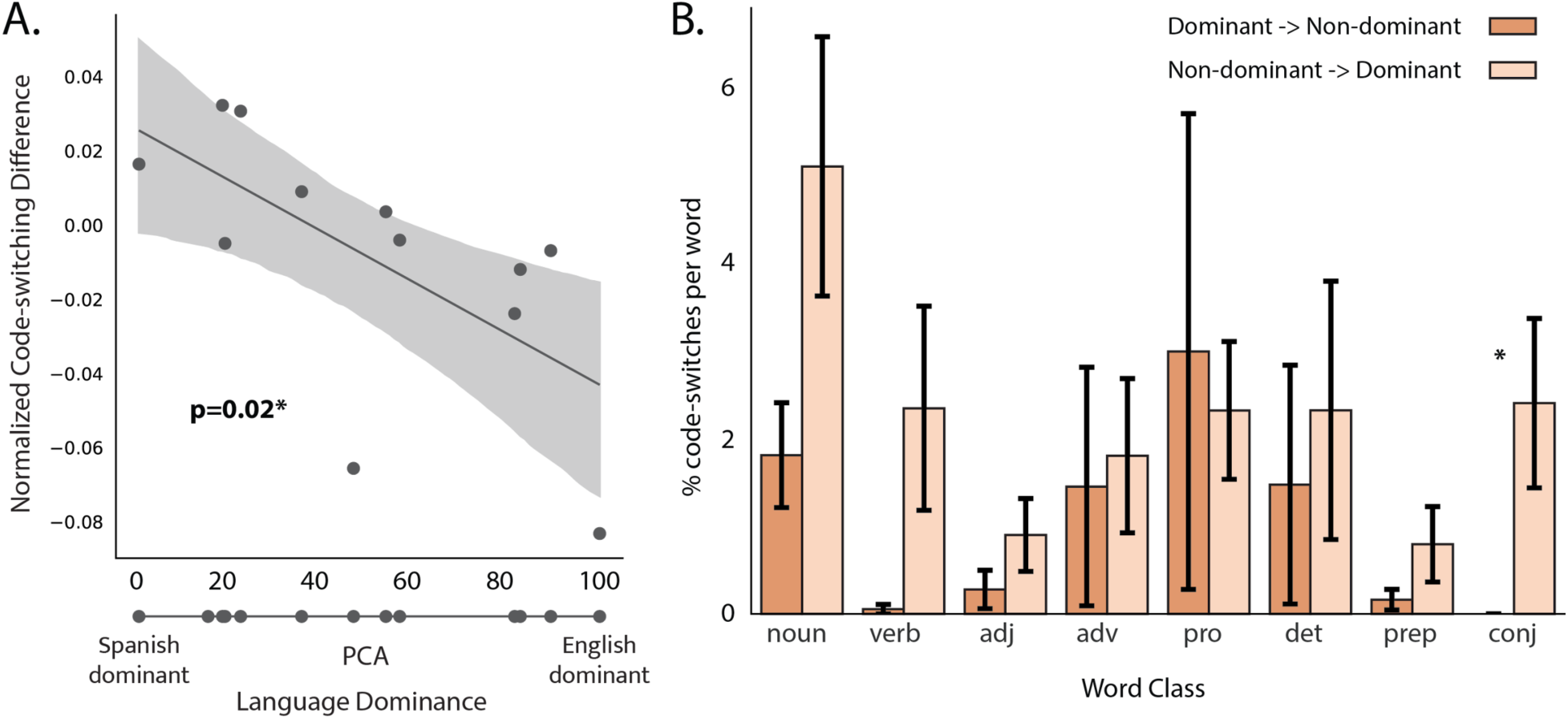
Effect of language dominance on the normalized code-switching measure. A: shows the correlation between PCA language dominance measure and the difference amount of code-switches. B: code-switching rate at each word class in the dominant and non-dominant language.

## 4 Discussion

### 4.1 Implications for bilingual language organization

The present study aimed to clarify the neurocognitive architecture of bilingual language representation by analyzing error patterns and code-switching behavior in Spanish-English bilingual individuals with post-stroke aphasia (PWA). We examined how different levels of representation (phonology, morphology, syntax, and lexico-semantics) behave in bilingual language breakdown and recovery, recognizing that convergence and dissociation can co-exist within a unified yet flexible system within the context of aphasia (Abutalebi & Green, 2013; Blanco-Elorrieta & Caramazza, 2021; Van de Putte et al., 2018).

A major strength of this study lies in the use of naturalistic, open-ended speech production tasks, which allowed us to evaluate bilingual language performance across all linguistic levels and word classes in ecologically valid discourse. Unlike tightly constrained experimental paradigms, narrative production affords the opportunity to capture spontaneous linguistic behavior as it unfolds in real-time, including morphological and syntactic structures that are often excluded from item-based assessments. This approach enabled us to conduct a fine-grained analysis of error types across phonology, morphology, syntax, and lexico-semantics, levels that are rarely examined simultaneously in bilingual aphasia research.

Strikingly, across participants, we found consistent qualitative profiles of impairment in both the dominant and non-dominant language not only at every level of representation but also across different word classes. This convergence across both structural levels and lexical categories provides especially robust evidence for a shared representational system. If the two languages were supported by partially or fully distinct mechanisms, some degree of divergence would be expected in at least one linguistic domain or lexical category. Yet even when broken down into distinct word types such as content vs. function words; nouns vs. verbs the same error typologies emerged in both languages, suggesting that the underlying representations and processing principles are largely shared across a bilingual’s two languages, even after brain damage, although quantitative differences in performance may emerge depending on pre-morbid proficiency and usage patterns. This pattern aligns with theoretical frameworks proposing that bilinguals rely on a shared representational system (e.g., Blanco-Elorrieta & Caramazza, 2021). Under this view, when bilinguals experience language breakdown, the impairment reflects damage to a single integrated system that supports both languages, rather than to two independent language-specific systems. Consistent with this, we found that the parallelism of error types across languages persisted even after controlling for language dominance and error class, and was independently confirmed through supervised classification analysis and unsupervised clustering analysis. Although some case reports have documented dissociations across languages (e.g., Paradis, 2001; Fabbro, 2001), our findings complement rather than contradict those observations. Such dissociations may arise in single cases lesion-specific factors or from differences in experiential variables such as age of acquisition and proficiency asymmetries, which can differentially strengthen components of the shared system. At the group level, our results indicate that shared representations constitute the default organizational principle of the bilingual language system, with divergence emerging only under specific individual conditions. Crucially, the presence of individual dissociations does not challenge integrated models if those dissociations reflect uneven disruption or use-based reinforcement within a common representational architecture, rather than separate neural substrates for each language.

We also observed clear quantitative differences, with more errors in the non-dominant language. These effects are consistent with frequency-based models of language access (Martin et al., 1996; Jacquemot et al., 2007; Gollan, 2014), and suggest that availability rather than distinct representational formats is the primary source of asymmetry. Future studies combining behavioral data with lesion localization will be important for disambiguating these possibilities.

Turning to code-switching, we found that PWAs switched significantly more from the non-dominant to the dominant language. This pattern is consistent with activation-based models of lexical selection (Blanco-Elorrieta & Caramazza, 2021) and contrasts with inhibitory models that predict more switching into the non-dominant language (Green & Abutalebi, 2013; Gollan et al., 2014). Importantly, we do not claim that our data refute all inhibitory models; rather, we propose that under naturalistic, self-paced conditions, lexical selection in bilingual PWAs appears to follow patterns of relative availability, not inhibition.

Prior studies documenting reverse dominance effects often rely on forced-switch or mixed-language paradigms (Li & Gollan, 2018), which engage domain-general control systems in a way that narrative production does not (Fedorenko & Blank, 2020). Our results instead align with recent behavioral findings in typical bilinguals and clinical populations that emphasize the communicative and facilitative role of switching (Goral et al., 2019; Bihovsky et al., 2023; Mooijman et al., 2024).

We also found a positive correlation between language proficiency and the extent of switching into the dominant language, further supporting the idea that such switches reflect facilitation. The study lacks a bilingual control group, which limits our ability to infer the aphasia-specific impact of these effects. This limitation is unavoidable, as neurologically healthy bilinguals would be unlikely to produce meaningful errors on the relatively simple set of AphasiaBank narrative tasks used in this study. Despite this constraint, our results implicate activation-based mechanisms as the primary force shaping bilingual lexical selection under impairment.

### 4.2 Clinical implications for bilingual aphasia

The present findings support a model in which bilingual individuals with aphasia share underlying linguistic representations across languages, even in the presence of quantitative asymmetries due to dominance. Given that neural damage typically disrupts (but does not fundamentally restructure) pre-existing language systems, our results also imply that these shared representations were likely present prior to stroke onset. Thus, while our data come from a clinical population, they lend indirect support to models of bilingualism that posit an integrated neurocognitive architecture in neurotypical individuals (e.g., Blanco-Elorrieta & Caramazza, 2021), that future research should test directly.

Clinically, this supports treatment approaches that target language-general mechanisms to promote cross-language transfer (Kiran & Roberts, 2010; Scimeca et al., 2024). The observed parallelism in error types across languages suggests that interventions targeting shared semantic or syntactic structures in one language may facilitate improvements in the other (Marte et al., 2025).

Code-switching behavior also has implications for diagnosis and therapy. Rather than interpreting code-switches as inherently pathological, our results indicate that bilingual PWAs may use switching as a strategy to maintain communicative effectiveness, consistent with the notion that switching can be compensatory rather than disruptive (Blanco-Elorrieta & Pylkkänen, 2017; Mooijman et al., 2024). While the current study did not directly measure communicative success, we interpret switching as an adaptive behavior likely grounded in activation-based selection mechanisms. On this basis, clinicians may benefit from considering assessment protocols that allow for responses in either language and permit switching where appropriate (Goral et al., 2024). Importantly, this interpretation should not be taken as a clinical prescription. Instead, we propose that clinicians consider communicative context and underlying intent when evaluating switching behavior.

### 4.3 Limitations and Future Directions

While our sample size is modest, it reflects the recruitment challenges inherent to this population. Excluding participants with limited discourse output introduces selection bias, as these individuals likely had more severe aphasia and/or significant disparities in proficiency. This exclusion may limit the generalizability of our findings to the full spectrum of bilingual aphasia severity. Longitudinal studies and larger samples are essential for evaluating whether recovery trajectories differ across levels of representation and between languages. Such work will be critical for refining theories of bilingual language organization and informing personalized, evidence-based interventions.

## 5 Conclusion

This study provides empirical support for a model of bilingual language organization in aphasia that relies on shared neural and cognitive resources across languages, modulated by relative language dominance. Parallel qualitative patterns of impairment across languages, together with activation-based code-switching profiles, suggest that bilinguals rely on an integrated language system that spans multiple levels of representation. These findings have both theoretical and clinical implications, advocating for models of bilingualism that incorporate shared architecture alongside usage-dependent variation, and for intervention strategies that leverage cross-language commonalities to support recovery in bilingual aphasia. Because neural injury reveals rather than creates underlying structures, these findings also support the inference that shared representations were present in the neurotypical brain prior to injury. However, direct comparisons with healthy bilinguals will be critical to fully establish which aspects of this integration are universal and which may be shaped by post-stroke adaptation.

## Data statement

The data will be publicly available in a OSF repository.

